# A reduced level of consciousness affects non-conscious processes

**DOI:** 10.1101/2020.11.10.376483

**Authors:** A. Fontan, L. Lindgren, T. Pedale, C. Brorsson, F. Bergström, J. Eriksson

## Abstract

Being conscious is a profound aspect of human existence, and understanding its function and its inception is considered one of the truly grand scientific challenges. However, the nature of consciousness remains enigmatic, to a large part because “being conscious” can refer to both the content (phenomenology) and the level (arousal) of consciousness, and how these different aspects are related remains unclear. To empirically assess the relation between level and content of consciousness, we manipulated these two aspects by presenting stimuli consciously or non-consciously and by using Propofol sedation, while brain activity was measured using fMRI. We observed that sedation greatly affected non-conscious processes, which starkly contrasts the notion that anesthetics selectively reduce consciousness. Our findings reveal that level and content of consciousness are separate phenomena, and imply that one may need to reconsider what “being conscious” means.

## Introduction

The concept of consciousness is multifaceted and can refer to at least two aspects: the content and the level/state of consciousness. The “content” relates to the core characteristic of consciousness, which is the subjective, phenomenal, “what-it-is-like” quality associated with experiencing something (Nagel, 1974). The level of consciousness commonly refers to arousal/wakefulness, and occurs on a continuum e.g., from comatose to fully awake (Laureys, 2005). These two aspects have mostly been investigated separately and there is much debate on how to conceptualize their relation (Bachmann, 2012; Bayne et al., 2016; Fazekas and Overgaard, 2016; Hohwy, 2009; Koch et al., 2016; Laureys, 2005; Overgaard et al., 2006).

On one hand, they can be considered as two aspects of the same underlying phenomenon (Aru et al., 2020; Bachmann and Hudetz, 2014; Mashour and Hudetz, 2017; Phillips et al., 2018; Suzuki and Larkum, 2020), which is supported by the observation that a certain level of arousal is required to enable conscious experiences. Indeed, we have a rich repertoire of conscious experiences when we are awake, and these experiences end during dreamless sleep or when we otherwise “lose” consciousness (Searle, 2000). In addition, brain research has demonstrated that the level of arousal affects integration of information across multiple brain regions (Casali et al., 2013), which may be key for generating conscious experiences (Tononi et al., 2016). Moreover, it has been suggested that neural mechanisms related to changes in the level of arousal overlap with the mechanisms generating conscious experiences (Aru et al., 2019), and general anesthesia is commonly considered to “selectively reduce consciousness”. Yet, while general anesthesia, sleep, or coma, are commonly described as states altering consciousness, the extent to which conscious experiences are lost/reduced when we are unresponsive is difficult to establish (Aru et al., 2020; Alkire et al., 2018; Bayne 2019).

On the other hand, the content and level/state of consciousness may be seen as separate phenomena (e.g., Bayne et al., 2016). A distinction between the two is apparent in every-day and clinical situations, which suggests instead that the level and the content of consciousness are not specifically interrelated. For example, vegetative-state patients can display sleep-wake cycles but remain unresponsive to external stimuli (Wislowska et al., 2017), and on rare occasions fully anesthetized patients can have conscious experiences (Errando et al., 2008). Moreover, we process information both consciously and non-consciously when we are awake (Kihlstrom, 1987).

To understand how level (hereafter referred to as “arousal”) and content (hereafter referred to as “conscious perception”) of consciousness are related, we set out to empirically assess their relation by manipulating both aspects while brain activity was measured using fMRI. Arousal was manipulated by administering two levels of the sedative Propofol. Importantly, participants were only mildly sedated and able to report whether they consciously perceived stimuli or not and to perform tasks during both sedation levels. Within each sedation level, the content of consciousness was manipulated by presenting visuospatial stimuli both consciously and non-consciously. Two possible outcomes may be expected. If reduced arousal selectively reduces processing of consciously perceived stimuli, the neural processes related to conscious perception would be uniquely affected by a change in arousal compared to non-conscious perception. Alternatively, neural processes would be affected by a change in arousal regardless of whether stimuli are consciously perceived or not.

## Results

This study included 30 healthy individuals who during fMRI performed a simple visuospatial main task under two levels of Propofol sedation: low (0.1 mg/h/kg; hereafter labelled “LS” for “low sedation”) and moderate sedation (“MS”). The visuospatial task was divided in blocks performed twice for each sedation level. A stabilization period (~6 min) was implemented between each block to allow the effect of Propofol to reach its steady state, during which participants performed a simple visuo-motor “metronome” task, consisting of timing their motor responses as synchronous as possible to a gray disc presented in one quadrant of the display. The main task, performed during stable periods of Propofol infusion, consisted of noting the location of a gray disc presented in one of the display’s quadrants. There were three presentation conditions: a conscious, a non-conscious, and an “absent” condition. Conscious/non-conscious perception was manipulated with continuous flash suppression (Tsuchiya and Koch, 2005) (Supplementary Fig. 1). After each trial, participants evaluated their visual experience of the disc on a three-point “perceptual awareness scale” (PAS; see Methods).

### Sedation effect on behavior

First, to ensure that participants’ arousal was affected by Propofol sedation, we verified that the response variability relative to the metronome response cue (i.e., how precisely participants paced the responses; Supplementary Fig.2A) increased for MS. Among the three stabilization periods, we observed that participants’ performance changed as a function of the Propofol level (F_2,58_ = 12.1; p = 0.0004). Indeed, variability increased with the change of sedation from LS to MS and decreased from MS to LS (Supplementary Fig.2B). This confirmed that participants’ arousal changed before each block of the main task.

During the main visuospatial task, comparison of PAS responses between the sedation levels revealed a significant interaction effect in conscious (F_2,58_ = 5.2, p = 0.009) and in non-conscious (F_2,58_ = 8.5, p = 0.0006) conditions. Specifically, the number of stimuli reported as unseen (PAS = 1) increased for MS relative to LS (Newman-Keuls test: p = 0.04 and p = 0.01 for conscious and non-conscious trials respectively), with a concomitant decrease of clear (PAS = 3) visual experiences in conscious (p = 0.02) and of vague (PAS = 2) visual experiences in non-conscious (p = 0.003) conditions. To ensure no conscious visual experience in non-conscious trials and clear perception in conscious trials, only trials with PAS = 1 in non-conscious and in absent conditions, and trials with PAS = 3 in conscious condition, were included (> 80% of trials in each condition for the two sedation levels; see Methods) in the following analyses.

For conscious trials, participants had near perfect accuracy (hits – false alarms; mean ± SD: LS = 0.99 ± 0.02; MS = 0.99 ± 0.04; Fig.1A), with no difference between sedation levels (Wilcoxon match pairs test: z = 0.27, p = 0.79). For non-conscious trials, accuracy was at chance level (mean ± SD: LS = 0.01 ± 0.12, t_30_ = 0.59; p = 0.56; MS = 0.006 ± 0.12; t_30_ = 0.25; p = 0.80; Fig.1A), again with no difference between sedation levels (Wilcoxon match pairs test: z = 0.20, p = 0.84). As such, these stimuli were non-conscious according to both subjective and objective criteria. Participants’ response time did not differ between non-conscious and absent trials for either sedation level (LS: t_29_ = −0.70, p = 0.5; MS: t_29_ = −0.32, p = 0.7), but was generally slower during MS compared to LS (main effect of sedation: F_1,29_ = 13.42; p = 0.0006; Fig.1B).

**Fig. 1:**
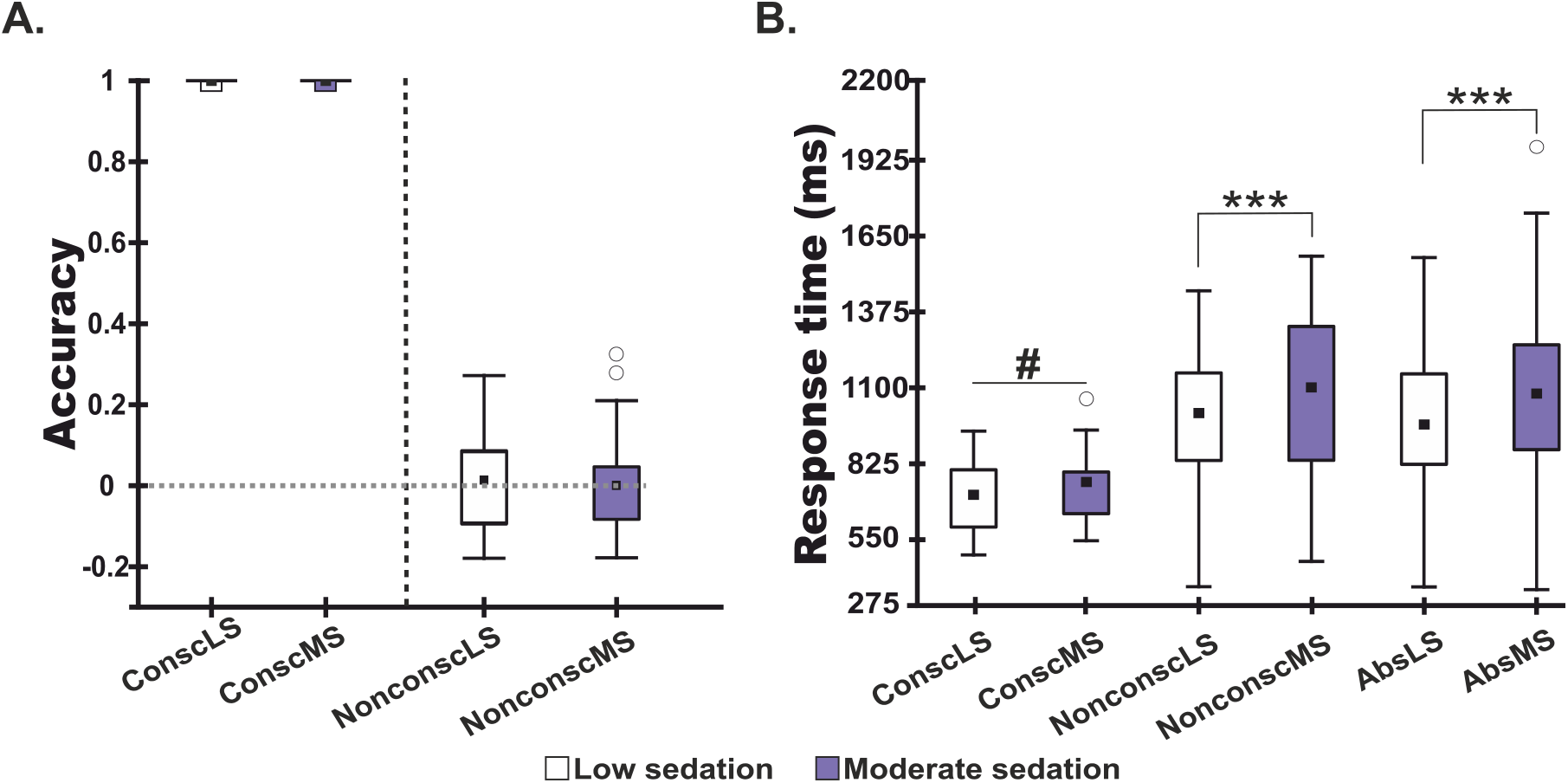
Behavioral performance during the visuospatial task under the two levels of sedation. **A.** Boxplots showing participants’ accuracy (hits – false alarms) to detect the correct stimulus location during low (white) and moderate (purple) sedation; the gray dashed line indicates chance level. **B.** A global slowing in response time verified that participants’ arousal was reduced with increased sedation. White circles indicate outliers; the difference was significant also without these data points. ***p< 0.001; # p=0.06.

### Neural response to stimulus presence

We then investigated the neural response related to both conscious and non-conscious visuospatial processing. Whole-brain univariate analyses of fMRI data, contrasting conscious to absent conditions, revealed significant blood-oxygen-level-dependent (BOLD) signal change in brain areas consistent with visuospatial processing (De Schotten et al., 2005) (Supplementary Fig.3). However, these analyses were not sensitive enough to reveal any signal change related to sedation levels, despite changes in participants’ response time during the main task and response variability during the metronome task. Non-conscious processing was not detected either. Multivariate pattern analysis (MVPA) is more sensitive compared to univariate analyses (Haxby, 2012), and has previously been used to investigate non-conscious processes (Ahrens, 2013; Bergström and Eriksson, 2018; Sheikh et al., 2019; Soto et al., 2019); it therefore appears better suited to capture the expectedly subtle BOLD signal changes related to MS and, crucially for the question at hand, if MS affects conscious and non-conscious processing differently.

Using MVPA, we first applied a searchlight approach to generate decoding accuracy maps of the mere presence of the stimulus for non-conscious trials (i.e., non-conscious *vs.* absent trials, irrespective of sedation level) for each individual separately. This searchlight decoding was restricted to brain areas previously shown to be involved in visuospatial perception (Wang et al., 2015) and were thresholded at 50% decoding accuracy. Corresponding maps for conscious *vs.* absent trials were also generated. Maps derived from both non-conscious and conscious trials included bilateral early visual cortex, intraparietal sulcus, and frontal eye fields (Fig.2A), and were used to define regions of interest (ROIs) to quantify the effect of sedation on conscious and non-conscious perceptual processing. To ensure that regional differences would not confound any differences detected between the sedation effects on conscious and non-conscious processing, the ROIs were defined as the overlap between the decoding maps for conscious and non-conscious trials (Fig.2A).

**Fig. 2:**
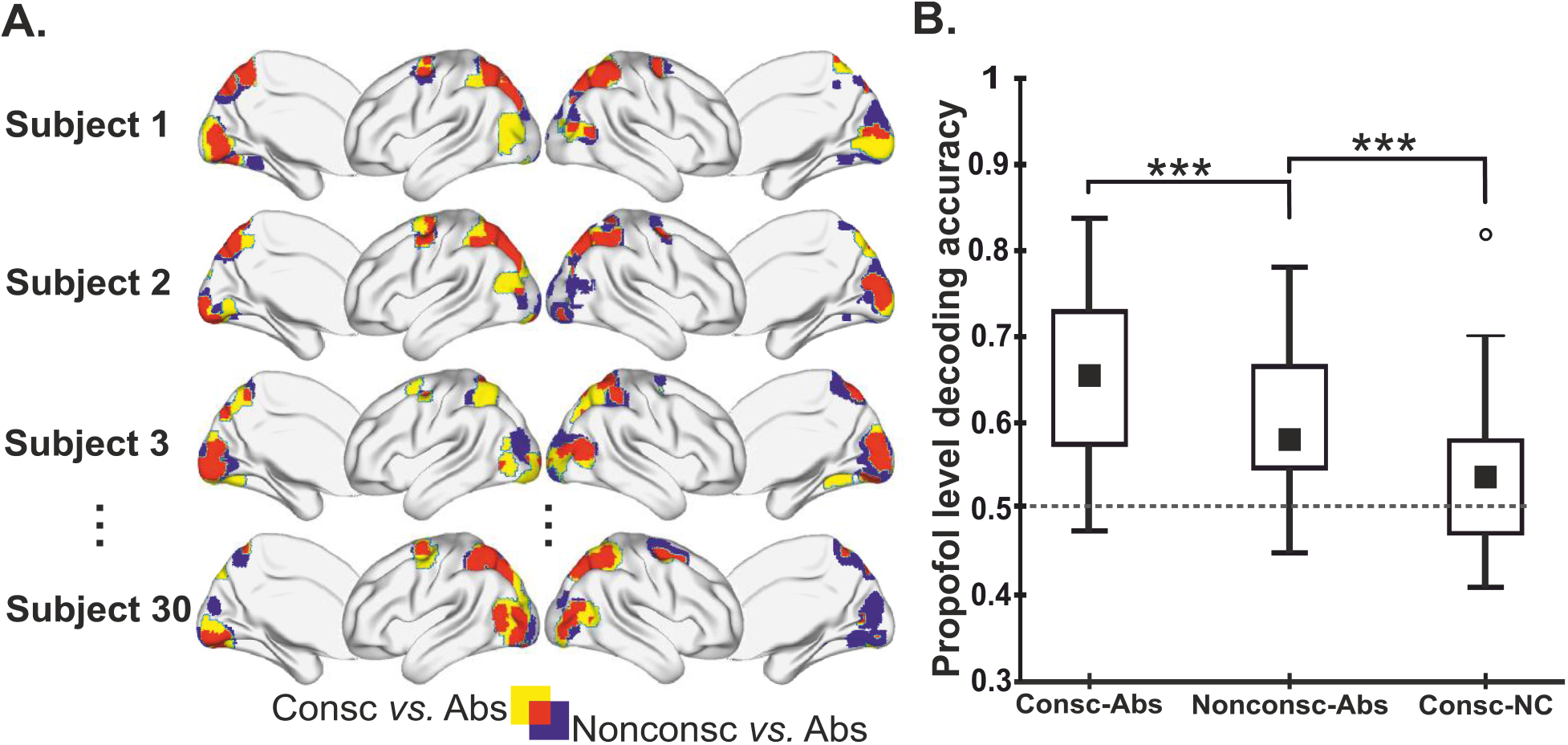
Effect of sedation on conscious and non-conscious visuospatial processing. **A.** Conscious (yellow) and non-conscious (blue) maps of MVPA searchlights, decoding presence *vs.* absence of the stimulus for each individual. Decoding of the sedation level was performed in a ROI defined as the overlap (red) between the two searchlight maps. **B.** Decoding accuracy for the sedation level classification (low *vs.* moderate), for conscious and non-conscious conditions controlled for unspecific sedation effects (-Abs), and for the conscious condition when controlling for non-conscious visuospatial processing (-NC). White circle indicates an outlier; the difference was significant also without this data points. Consc: Conscious; Nonconsc: non-conscious. ***p < 0.001.

### Sedation effect on neural patterns

We then used a ROI-based MVPA to decode the difference between LS and MS for conscious and non-conscious trials separately. In addition, we used a contrast approach on a trial-by-trial basis for the MVPA classification to control for possible non-specific effects of visuospatial processing. Thus, the BOLD signal from the absent condition was subtracted from conscious and non-conscious BOLD signal, separately for each sedation level (i.e., ConscLS-AbsLS vs ConscMS-AbsMS, NonconscLS-AbsLS vs NonconscMS-AbsMS; see Materials and Methods for more details). Possible non-specific changes in cerebral blood flow were also quantified using arterial spin labeling and no significant difference between LS and MS was observed (see Methods).

Decoding accuracies for the sedation effect on conscious processing (65.0 ± 10.2%) and on non-conscious processing (60.8 ± 8.5%) were significantly different from chance (permutation test (Stelzer et al., 2013): p = 0; i.e., all 10000 permutations had a value below 65.0 and 60.8%, respectively), revealing that the sedation affected both conscious and non-conscious processes. Moreover, we observed a significant difference between conscious and non-conscious decoding accuracies (Wilcoxon matched pairs test: z = 4.63; p = 3.5×10^−6^), revealing an interaction between arousal and conscious/non-conscious visuospatial processing, thereby demonstrating that a change in the arousal affects conscious and non-conscious processes differently.

This higher decoding accuracy for the sedation effect on conscious processing compared to non-conscious processing would thus indicate that arousal does indeed primarily affects conscious processing. Crucially, these results are based on BOLD signal changes where a common reference condition (absent trials) had been subtracted. While controlling for non-specific effects from Propofol sedation, this subtraction procedure does not isolate conscious processing from non-conscious processing. Arguably, BOLD signal for conscious trials should therefore partly reflect the non-conscious processes that precede and lead to conscious perception of the target stimulus, in addition to conscious processes. To ensure that non-conscious processes did not drive the decoding of sedation levels in the conscious condition, we performed a further analysis where non-conscious BOLD signal was subtracted from conscious BOLD signal, separately for each sedation level (i.e., ConscLS-NonsconscLS vs ConscMS-NonconscMS, see Materials and Methods for details). Surprisingly, while the decoding accuracy of the sedation level for conscious processing (54.8 ± 7.3%) was significantly different from chance (permutation test: p = 0.001), it was also significantly lower than the decoding accuracy of the sedation effects on non-conscious processing (Wilcoxon matched pairs test: z = 2.9; p = 0.0032), revealing that arousal affected non-conscious processes even more than it affected conscious processes (Fig.2B). These results were replicated in supplemental analyses using alternative ROIs based on the metronome task performed during the stabilization periods, which, similar to the main task, requires visuospatial processing (see Supplementary Fig.5).

## Discussion

The finding that a reduced level of arousal greatly affects non-conscious processes has several implications for the concept of consciousness and for clinical situations. Firstly, it could be argued that the conception of “levels of consciousness” when referring to arousal is a misnomer. While our results do show that conscious processes are affected by a reduced level of arousal, even greater changes were evident for non-conscious processes. Thus, to denote a reduced level of arousal as an altered level (or state) of consciousness is potentially misleading, because such terminology suggests a specificity that apparently is non-existent. It may be more appropriate to simply use “arousal”, “alertness”, or similar terminology, because a change in arousal affects both conscious and non-conscious processing, which together constitute the individual’s mental capacity. Our findings are consistent with theories of consciousness that explicitly separate levels and content (e.g., Northoff and Huang, 2017), but are problematic for hypotheses that suggest integrating these two dimensions as subtending consciousness (e.g., Aru et al., 2019; Bachmann and Hudetz, 2014).

Secondly, our findings have implications for the development of “consciousness markers” in people with low levels of arousal and/or altered “states of (un)consciousness”. One great challenge in consciousness research, which has substantial ethical implications, is to know whether patients that are non-responsive due to anesthesia or trauma retain their capacity for conscious experiences. One candidate marker, suggested to reflect conscious experience (i.e., content consciousness), is the perturbational complexity index (PCI), which reliably discriminate between lower “levels of consciousness”, including sleep, anesthesia, and in patients with consciousness disorders (Casali et al., 2013; Sarasso et al., 2015). To our knowledge, the PCI and other candidate markers do not take changes in non-conscious processing into account. While the sedation-related changes observed in conscious processes are consistent with the idea of reduced integration (Schrouff et al., 2011), we have here shown that changes in non-conscious processing were substantial. Again, the lack of specificity, i.e., the inability to isolate processing specifically related to (the content of) consciousness and to exclude effects emanating from changes in non-conscious processing, is problematic for existing markers of consciousness. The same goes for research on the neural correlates of consciousness where manipulations of arousal are used (Eriksson et al., 2020; Koch et al., 2016). That is, given our current findings, changes in markers/indices or correlates are likely to have been driven by non-conscious in addition to conscious processes. For practical purposes, e.g., when trying to determine if a patient is capable of conscious experiences or not, a correlation between general mental capacity and the capacity for conscious experiences may suffice, but should be verified.

Previous neuroimaging research on the effects of sedation has demonstrated a sparing of neural activity in sensory regions combined with a reduction in higher-order regions, including frontal and parietal cortex (e.g.,Demertzi et al., 2019; Hudetz and Mashour, 2016). Such previous findings are consistent with the notion that sedation primarily affects conscious perceptual processing, but are not necessary inconsistent with our current findings. Indeed, while the current study is the first to manipulate both arousal and conscious perception simultaneously, we have only used two levels of sedation. It therefore remains unknown whether the relation is linear or non-linear.In addition, the smaller impact of the MS on conscious processes, and that is also consistent with participants’ accuracy in this condition, could result from some form of attentional effect that acts differently on conscious vs. non-conscious stimuli (Coull et al., 2004). Further characterization of the effects of alertness on conscious and non-conscious processing is an important task for future research, together with investigations of how the current findings generalize to other stimuli, tasks, and manipulations of alertness.

In conclusion, our current results show that Propofol greatly alters non-conscious processes and to a lesser extent conscious processes, contrary to the notion that anesthetics selectively reduce “consciousness”. This finding implies that one may need to reconsider what it means to “be conscious”, and could lead to improved markers of consciousness and to a better understanding of situations where content and level dissociate, for example when sedated patients retain the capacity for conscious experiences.

## Materials and Methods

### Participants

Forty healthy right-handed adults took part in the experiment. Participants were recruited from Umeå University campus through poster and internet advertisements. They had normal or corrected-to-normal vision, right-eye dominance, gave their written informed consent, and received financial compensation for participation (600 SEK). Ten participants were excluded from the analyses, either due to excessive head movement during fMRI scanning (n = 5), for failing to follow task instructions (n = 4), or because non-conscious processing could not be verified (n = 1). Thus, the final sample in the analyses was 30 individuals (mean age ± SD: 27.4 ± 4.6 years; 12 males). This study was approved by the regional ethics review board (dnr 2018-314-32M).

### Paradigm and stimuli

During fMRI scanning, participants performed a visuospatial task, under continuous flash suppression (CFS), composed of 120 trials equally distributed in 2 blocks and divided into 3 presentation conditions: 40 conscious, 60 non-conscious, and 20 absent trials for each sedation level. Each trial was randomly chosen from one of the three conditions.

For CFS, a mirror stereoscope was used to isolate visual input from left and right side of the screen to participants’ corresponding eyes. For non-conscious trials, the target stimulus (gray disc; size = 0.6°) was presented for 500 ms to the non-dominant (left) eye while colored squares of random composition (“Mondrians”; size = 4.2° × 4.2°) where flashed (10 Hz) to the dominant eye to suppress conscious experience of the disc (Tsuchiya and Koch, 2005). Mondrians were flashed for 500 ms longer than the disc’s presentation, minimizing the risk of adaptation after - effects. To maximize stimulus intensity during non-conscious trials, contrast between the disc and the gray background was increased or decreased every 10 trials depending on how many times participant reported the disc as seen. That way, the proportion of actual non-consciously experienced disc presentations was 80%. There were 17 possible contrast values. The difference between each contrast consisted of an increase or a decrease in RGB value of 2 (range = 174-206; background = 210). For conscious trials, the disc (RGB = 198) was superimposed on Mondrians, presented to the dominant eye, and was thus consciously seen. For “Absent” trials, used as reference condition, Mondrians were presented to the dominant eye while an empty gray background (4.2° × 4.2°) was presented to the non-dominant eye.

For conscious and non-conscious trials, the disc was presented in one of the four quadrants of the screen. The position was randomly selected from a pre-specified list where positions were counterbalanced within each condition. After the disc presentation, a probe was presented, pointing either to the same spatial location as the disc (match) or to another spatial location (non-match). Participants had to decide whether the probe was pointing to the disc’s location (yes/no). For non-conscious and absent trials, participants were instructed to guess on the first alternative that came to mind. There was 50% chance that the probe pointed to the disc location. After the probe, participants estimated their conscious experience of the disc on a three-point perceptual awareness scale (PAS) (Sandberg et al., 2014), from 1: no visual experience to 3: clear visual experience of the disc. For probe and PAS, participants had to reply within a limit of 2.5 s after which the experiment automatically continued to the next trial. The inter-trial interval (ITI) was adjusted according to participants’ response time in a way that two trials were always separated by 5 s.

A visual metronome task was also performed and used as a behavioral measure of participants’ arousal. Participants had to synchronize finger taps to visual isochronous metronome sequences presented to their dominant eye, and were requested not to follow the beat by moving other body’s parts or using covert counting. The stimulus was the same disc as for the visuospatial task but presented on the gray background with empty dotted circles reflecting the 4 possible positions of the stimulus apparition. One trial consisted of a 500-ms stimulus presentation followed by a 500-ms ITI. In total, participants completed 12 sequences of 20 trials (240 visual presentations) where the stimulus was presented with a 1-Hz tempo. The stimulus’ position within each block was selected in a pseudo-random order in a way that a block mainly consisted (85% of the trials) of stimuli appearing in one quadrant. Finally, participants received feedbacks about their performance at the end of each sequence.

### Propofol sedation: individual adjustment

The anesthetic agent used to manipulate arousal was Propofol (20 mg/ml), which activates GABAa receptors directly (O’Shea et al., 2000). Propofol is considered safe and fast acting (reaches its steady state ~6 min after infusion) (Trapani et al., 2012), which allowed us to change the level of arousal several times during fMRI session. Here, two sedation levels were used: a moderate level that was adjusted individually, and a low level (0.1 mg/kg/h). The choice of having 0.1 mg/kg/h rather than no sedative or saline injection, as a state of comparison was motivated by the fact that the sedative may affect blood flow or other non-neuronal parameters relevant to the fMRI signal (Qiu et al., 2017).

Individual adjustment of the moderate sedation level was evaluated during a pre-scanning session. Participants fasted from solids for at least 6h and from liquids 4h before sedation. Propofol was infused through an intravenous catheter placed into a forearm vein. Sedation was achieved using computer-controlled intravenous infusion of Propofol to obtain constant effect-site concentrations. Participants where initially injected with 2.0 mg/kg/h of Propofol. The infusion rate was then increased in steps of 0.25 mg/kg/h, separated by a 6-min stabilization period, until participants were considered moderately sedated, operationalized as when they showed difficulties to keep their eyes open, but remained responsive. Physiological parameters such as blood pressure, pulse oximetry, and breathing frequency were continuously monitored and were stable during Propofol infusion, and no side effects were observed. Anesthesia administration and monitoring were based on clinical judgment of the anesthesiologist and the intensive care nurse. In the final population (n = 30), the range of the moderate sedation was 2.25 to 4.0 mg/kg/h (mean ± SD: 2.8 ± 0.5 mg/kg/h).

### MRI data collection

Propofol infusion started right before participants were placed in the scanner bore. Participants began the experiment with either the low (“LS”) or the moderate (“MS”) sedation. In the final sample, 17 participants started with LS and 13 with MS. A certified intensive-care nurse with specific responsibility for pharmacological administration and monitoring was present throughout the session, and complete resuscitation equipment was available at all times.

The session started with structural imaging (T1, T2 FLAIR and T2 PROPELLER sequences) so that Propofol levels could stabilize before fMRI scanning. Then, one resting-state fMRI sequence was run at each sedation level for the use of another study and will not be further reported here, and task fMRI followed.

During task-fMRI, participants performed two 7-min blocks of the visuospatial task under both sedation levels. Each block was followed by a 6-min stabilization period where sedation level was changed and during which participants performed the visual metronome task. This resulted in 4 blocks of visuospatial task and 3 stabilization periods. Finally, to verify that Propofol was not interfering with regional cerebral blood flow (CBF) response at the sedative concentrations (Veselis et al., 2005), and did not modify flow-metabolism coupling (Johnston et al., 2003), the MRI session included two pulsed arterial spin-labeling (ASL) sequences for each sedation level.

MRI data were collected with a General Electric 3 Tesla Discovery MR750 scanner (32-channel receive-only head coil). High-resolution T1-weighted structural image was collected FSPGR with TE = 3.2 ms, TR = 8.2 ms, TI = 450 ms, and flip angle = 12°. Task-fMRI (1410 volumes) was recorded using a T2*-weighted gradient echo pulse sequence, echo planar imaging, field of view = 25 cm, matrix size = 96 × 96, slice thickness = 3.4 mm. The volumes covered the whole cerebrum and most of the cerebellum containing 37 slices with 0.5 mm inter-slice gap and an ASSET acceleration factor of 2. The orientation was oblique axial, and slices were aligned with the anterior/posterior commissures, and scanned in interleaved order with TE = 30 ms, TR = 2 s, flip angle = 80°.

Finally, ASL was collected using a field of view = 24 cm, matrix size = 128 x 128, bandwidth of 62.50 kHz; slice thickness = 4 mm. The acquisition orientation was axial aligned with the anterior/posterior commissures. The 40 slices with 2 mm inter-slice spacing were acquired from inferior to superior in an interleaved order to cover most of the cortex with a TR = 4 s.

### Data processing and statistical analyses

In the visuospatial task, trials with response time (RT) < 250 ms or > 2.5 s were excluded prior to statistical analyses (Ratcliff, 1993). Then, PAS responses between LS and MS during conscious or non-conscious trials were compared using repeated-measure two-way analysis of variance (ANOVA). Afterwards, only trials in absent (LS: 87.6 ± 15.9 %; MS: 87.5 ± 14.7 % of trials) and non-conscious (LS: 81.3 ± 18.1 %; MS: 85.1 ± 18.7 % of trials) conditions with PAS = 1, and trials with PAS = 3 in conscious (LS: 97.5 ± 3.0 %; MS: 94.3 ± 7.0 %) condition were included in the analyses.

For the accuracy analyses, a hit was defined as a position match between disc location and probe together with a “yes” response, while a “no” response was defined as a miss. False alarm (FA) was considered as a non-match between disc location and probe with a “yes” response, while a “no” response defined a correct rejection (CR). Accuracy was defined as the proportion of correct answers (hits-FA) for conscious and non-conscious trials.

Accuracy, under the two sedation levels, was compared using Wilcoxon’s matched pairs test in conscious and in non-conscious conditions. RT differences between the two sedation levels were assessed using repeated-measure two-way ANOVA across the three visual presentation conditions. Specific differences for RT in MS and in LS between non-conscious and absent conditions were evaluated using Student’s t-tests.

For the metronome task, the three first trials of each sequence and missed responses were discarded from analysis to include only trials where participants were synchronized to the stimulus.

Visual-to-tap asynchrony was calculated as the absolute time difference between stimulus onset and participant’s response. In other words, the smaller the difference, the better the performance. Then, variability in asynchrony was calculated for each sequence and each participant. Changes in variability due to Propofol sedation were estimated with the slope of a linear regression across the 12 sequences, and were used as a sedation-effect estimation. A positive slope (increased variability) with increased Propofol reflected a decrease in arousal and vice versa. To assess changes in the sedation effect over the three stabilization periods at the group level, the sign of the slopes related to participants who started the experiment with MS was switched, respectively for each stabilization period. Group level comparison was done using repeated-measure one-way ANOVA.

All post hoc tests with correction for multiple comparisons were conducted using Newman-Keuls test and p-value < 0.05 was considered significant.

### fMRI analyses

Image pre-processing, statistical fMRI, and ASL data analyses were conducted with SPM12 (Wellcome Department of Imaging Neuroscience, London, UK) running in Matlab 8.4 (Mathworks, Inc., Sherbon, MA, USA) using custom-made Matlab scripts. Functional images were (i) slice-time corrected, (ii) realigned to the first image of the time series to correct for head movement, (iii) unwarped to remove residual movement-related variance (Andersson et al., 2001), and (iv) co-registered to high-resolution structural data. Structural images were normalized to the MNI (Montreal Neurological Institute) template using DARTEL (Ashburner, 2007) and resulting parameters were used for functional images normalization, which were resampled to 2-mm isotropic voxel size. Finally, functional images were smoothed with an 8-mm and a 2-mm FWHM Gaussian kernel for univariate and multivariate pattern analysis (MVPA) (Gardumi et al., 2016) respectively.

### Univariate analysis

Pre-processed data were analyzed using a two-stage summary statistics random effect model (Friston et al., 1995; Holmes and Friston, 1998). At the first stage, task-dependent changes in BOLD signal were modeled as zero-duration event regressors time-locked to (i) the Mondrians’ onsets for the visuospatial task, including conscious, non-conscious and absent conditions for each Propofol level and each PAS rating, and to (ii) the stimulus’ onsets for the visual metronome task, including the four stimulus’ positions. These regressors were convolved with the SPM12 canonical hemodynamic response function and entered into general linear model (GLM). The models also included constant terms, 6 head movement parameters, nuisance regressors such as missed responses, and physiological noise (6 parameters) from white matter and cerebrospinal fluid, estimated using aCompCor method (Behzadi et al., 2007). Finally, high-pass filter (cut-off = 128 s) was applied to remove low-frequency drifts in the data.

Contrast maps were computed on beta maps resulting from the estimated first-level GLMs to reveal for conscious and non-conscious conditions, brain regions (i) subtending visuospatial processing regardless of sedation levels and (ii) presenting differences between sedation levels. Individuals’ maps subtending conscious and non-conscious visuospatial networks were taken to second-level random-effects analyses (one-sample t-tests) to account for inter-individual variability. Comparison between sedation levels was done using paired t-tests for conscious and non-conscious conditions.

For the ASL data, the mean CBF value for gray matter for both sedation levels was calculated using histogram-based segmentation algorithm of the upper brain CBF values, based on the ASL sequences. Averaged difference images were converted to mL/100g/min using a single-compartment model. CBF images were (i) co-registered to high-resolution structural data, (ii) motion-corrected using a 6-parameters rigid body spatial transformation, and (iii) normalized to the MNI via DARTEL template image. CBF images for each participant were taken to second-level random-effects analyses (paired t-tests) to estimate CBF differences as a function of Propofol level.

Multiple comparisons correction of statistical maps at the second level was conducted on the whole brain using cluster-based extent thresholding of p < 0.05 (FWE corrected) calculated based on the Gaussian random field method and following cluster-defining threshold of p < 0.001.

### MVPA: Defining regions of interest

Two searchlight MVPAs were conducted, for conscious and for non-conscious trials, to identify regions of interest (ROIs) where the mere presence of the stimulus could be decoded irrespective of sedation level (conscious *vs.* absent and non-conscious *vs.* absent). A searchlight decoding approach was used (Grootswagers et al., 2017; Haynes, 2015; Kriegeskorte et al., 2006; Pereira et al., 2009) as implemented in CoSMoMVPA decoding toolbox (Oosterhof et al., 2016), on the 2-mm smoothed beta parameter maps from the GLM described above (one map per trial). The number of maps/trials were balanced for each participant across conditions (Mumford et al., 2012). Then, a spherical searchlight (~300 voxels) was used to extract local features for classification, and was moved across the search space. To specifically identify regions subtending visuospatial perception, the search space was limited to probabilistic maps of visual topography (Wang et al., 2015). A linear discriminant analysis (LDA) classifier was used, combined with a 10-fold cross-validation procedure (Varoquaux et al., 2017). Within-run cross-validation has been shown to be unbiased for randomized event-related designs, as used here (Mumford et al., 2014). Nevertheless, to ensure that BOLD signal was non-overlapping between validation folds, we included only trials such that there was at least 30 s between the training and the testing fold. Individual maps of classification accuracies were thresholded at 50% (chance level) and smoothed with an 8-mm FWHM Gaussian kernel.

To ensure that the decoding of Propofol effects for conscious and non-conscious trials would be comparable and not confounded by regional differences, the ROI was defined such that both conscious and non-conscious processing was assuredly present within the ROI for each individual. That is, the overlap between the above-described searchlights was selected as ROI for each individual (ROI range size = 1168-2487 voxels). Importantly, the ROI-defining comparisons of conscious/non-conscious *vs.* absent are orthogonal to the latter comparisons of sedation levels (Kriegeskorte et al., 2009). Nevertheless, to verify that the above procedure for defining ROIs did not affect the latter classification of sedation levels, a third searchlight MVPA, using the metronome task data, was performed identifying again ROI of visuospatial processing. Specifically, the stimulus’ position was decoded (left vs. right). For comparison with the original ROIs, the same number of voxels, for each individual, was used in these alternative ROI analyses.

### MVPA: Decoding Propofol sedation

To quantify the sedation effect (in terms of decoding accuracy) specifically related to conscious and to non-conscious visuospatial processing, we first analyzed it in relation to a common baseline – the absent conditions, using a contrast on a trial-by-trial basis. Thus, beta maps from the absent conditions were subtracted from conscious (ConscLS-AbsLS and ConscMS-AbsMS) and from non-conscious (NonconscLS-AbsLS and NonconscMS-AbsMS) beta maps separately for each sedation level. To retain power although there were fewer absent trials than conscious/non-conscious trials, absent beta maps were randomly selected within each block such that the same beta map of the absent condition could be used no more than three times to be subtracted from conscious or from non-conscious beta maps. This level-specific subtracting procedure controls for non-specific effects from Propofol (e.g., subtle changes in CBF) and for effects unrelated to visuospatial processing.

Arguably, conscious perceptual experiences are preceded by non-conscious processing (Aru et al., 2019). Thus, to ensure that the sedation level decoding in the conscious condition was specifically related to conscious experiences, a second procedure of subtracting level-specific non-conscious beta maps was performed. Here, non-conscious beta maps were subtracted from conscious beta maps for LS and MS separately (ConscLS-NC and ConscMS-NC). Because there were more non-conscious than conscious trials, the surplus non-conscious trials were randomly selected for exclusion within each block.

Three ROI-based MVPAs, using the ROIs described above, were thus performed to decode the sedation level: (i) ConscLS-AbsLS *vs.* ConscMS-AbsMS, (ii) NonconscLS-AbsLS *vs.* NonconscMS-AbsMS, and (iii) ConscLS-NCLS *vs.* ConscMS-NCLS. Classification of Propofol levels was performed on a balanced number of beta maps/trials for each participant across conscious/non-conscious conditions and across the two sedations levels, thereby ensuring that decoding accuracy would be comparable and not confounded by the number of trials used in the classification. A LDA classifier and 10-fold cross-validation with at least 30 s between the testing and the training folds was used.

Individual accuracy values were entered into second-level analyses, using first a two-step permutation procedure (Stelzer et al., 2013) to evaluate if classification at the group level was significantly above chance level. Finally, to evaluate whether MS affected non-conscious and conscious processing differently, differences in decoding accuracy between conscious and non-conscious were assessed using Wilcoxon’s matched pairs test. A p-value < 0.05 was considered significant.

## Acknowledgments

We thank Göran Westling for engineering support, Anders Lundquist for statistical advice, Anders Wåhlin for support with the analysis of ASL data, and Stefan Lehtipalo and Johan Söderberg for support during pilot studies. We also thank George Northoff, Christof Koch, and Lars Nyberg for comments on previous versions of the manuscript, and all staff at UFBI for their contribution. This study was supported by Riksbankens Jubileumsfond (Swedish foundation for humanities and social sciences; Grant: P17-0772:1). FB was supported by Fundação para a Ciência e Tecnologia (CEECIND/03661/2017).

## References

Ahrens M. 2013. Multivariate Pattern Analysis of fMRI Data for Functional Voice Localizer. Neuroimage 62:852–855. doi:10.1016/j.neuroimage.2012.03.016.Multivariate

Andersson JLR, Hutton C, Ashburner J, Turner R, Friston K. 2001. Modeling geometric deformations in EPI time series. Neuroimage 13:903–919. doi:10.1006/nimg.2001.0746

Aru J, Suzuki M, Larkum ME. 2020. Cellular Mechanisms of Conscious Processing. Trends Cogn Sci xx:1–12. doi:10.1016/j.tics.2020.07.006

Aru J, Suzuki M, Rutiku R, Larkum ME, Bachmann T. 2019. Coupling the State and Contents of Consciousness. Front Syst Neurosci 13:1–9. doi:10.3389/fnsys.2019.00043

Ashburner J. 2007. A fast diffeomorphic image registration algorithm. Neuroimage 38:95–113. doi:10.1016/j.neuroimage.2007.07.007

Bachmann T. 2012. How to begin to overcome the ambiguity present in differentiation between contents and levels of consciousness? Front Psychol 3:1–6. doi:10.3389/fpsyg.2012.00082

Bachmann T, Hudetz AG. 2014. It is time to combine the two main traditions in the research on the neural correlates of consciousness: C=LxD. Front Psychol 5:1–13. doi:10.3389/fpsyg.2014.00940

Bayne T, Hohwy J, Owen AM. 2016. Are There Levels of Consciousness? Trends Cogn Sci 20:405–413. doi:10.1016/j.tics.2016.03.009

Behzadi Y, Restom K, Liau J, Liu TT. 2007. A component based noise correction method (CompCor) for BOLD and perfusion based fMRI. Neuroimage 37:90–101. doi:10.1016/j.neuroimage.2007.04.042

Bergström F, Eriksson J. 2018. Neural evidence for non-conscious working memory. Cereb Cortex 28:3217–3228. doi:10.1093/cercor/bhx193

Casali AG, Gosseries O, Rosanova M, Boly M, Sarasso S, Casali KR, Casarotto S, Bruno MA, Laureys S, Tononi G, Massimini M. 2013. A theoretically based index of consciousness independent of sensory processing and behavior. Sci Transl Med 5:198ra105. doi:10.1126/scitranslmed.3006294

Coull JT, Jones MEP, Egan TD, Frith CD, Maze M. 2004. Attentional effects of noradrenaline vary with arousal level: Selective activation of thalamic pulvinar in humans. Neuroimage 22:315–322. doi:10.1016/j.neuroimage.2003.12.022

De Schotten MT, Urbanski M, Duffau H, Volle E, Lévy R, Dubois B, Bartolomeo P. 2005. Neuroscience: Direct evidence for a parietal-frontal pathway subserving spatial awareness in humans. Science (80-) 309:2226–2228. doi:10.1126/science.1116251

Demertzi A, Tagliazucchi E, Dehaene S, Deco G, Barttfeld P, Raimondo F, Martial C, Fernández-Espejo D, Rohaut B, Voss HU, Schiff ND, Owen AM, Laureys S, Naccache L, Sitt JD. 2019. Human consciousness is supported by dynamic complex patterns of brain signal coordination. Sci Adv 5:1–12. doi:10.1126/sciadv.aat7603

Eriksson J, Fontan A, Pedale T. 2020. Make the Unconscious Explicit to Boost the Science of Consciousness. Front Psychol 11:1–4. doi:10.3389/fpsyg.2020.00260

Errando CL, Sigl JC, Robles M, Calabuig E, García J, Arocas F, Higueras R, Del Rosario E, López D, Peiró CM, Soriano JL, Chaves S, Gil F, García-Aguado R. 2008. Awareness with recall during general anaesthesia: A prospective observational evaluation of 4001 patients. Br J Anaesth 101:178–185. doi:10.1093/bja/aen144

Fazekas P, Overgaard M. 2016. Multidimensional Models of Degrees and Levels of Consciousness. Trends Cogn Sci 20:715–716. doi:10.1016/j.tics.2016.06.011

Friston KJ, Holmes AP, Poline JB, Grasby PJ, Williams SCR, Frackowiak RSJ, Turner R. 1995. Analysis of fMRI time-series revisited. Neuroimage. doi:10.1006/nimg.1995.1007

Gardumi A, Ivanov D, Hausfeld L, Valente G, Formisano E, Uludağ K. 2016. The effect of spatial resolution on decoding accuracy in fMRI multivariate pattern analysis. Neuroimage 132:32–42. doi:10.1016/j.neuroimage.2016.02.033

Grootswagers T, Wardle SG, Carlson TA. 2017. Decoding dynamic brain patterns from evoked responses: A tutorial on multivariate pattern analysis applied to time series neuroimaging data. J Cogn Neurosci 29:677–697. doi:10.1162/jocn_a_01068

Haxby J V. 2012. Multivariate pattern analysis of fMRI: The early beginnings. Neuroimage 62:852–855. doi:10.1016/j.neuroimage.2012.03.016

Haynes JD. 2015. A Primer on Pattern-Based Approaches to fMRI: Principles, Pitfalls, and Perspectives. Neuron 87:257–270. doi:10.1016/j.neuron.2015.05.025

Hohwy J. 2009. The neural correlates of consciousness: New experimental approaches needed? Conscious Cogn 18:428–438. doi:10.1016/j.concog.2009.02.006

Holmes AP, Friston KJ. 1998. Generalisability, random effects & population inference. Neuroimage 7:54–57. doi:10.1016/s1053-8119(18)31587-8

Hudetz AG, Mashour GA. 2016. Disconnecting Consciousness: Is There a Common Anesthetic End Point? Anesth Analg 123:1228–1240. doi:10.1213/ANE.0000000000001353

Johnston AJ, Steiner LA, Chatfield DA, Coleman MR, Coles JP, Al-Rawi PG, Menon DK, Gupta AK. 2003. Effects of propofol on cerebral oxygenation and metabolism after head injury. Br J Anaesth 91:781–786. doi:10.1093/bja/aeg256

Kihlstrom JF. 1987. Unconscious The Information-Processing Perspective. Science (80-) 237:1445–1452.

Koch C, Massimini M, Boly M, Tononi G. 2016. Neural correlates of consciousness: Progress and problems. Nat Rev Neurosci 17:307–321. doi:10.1038/nrn.2016.22

Kriegeskorte N, Goebel R, Bandettini P. 2006. Information-based functional brain mapping. Proc Natl Acad Sci U S A 103:3863–3868. doi:10.1073/pnas.0600244103

Kriegeskorte N, Simmons WK, Bellgowan PSF, Baker CI. 2009. Circular analysis in systems neuroscience – the dangers of double dipping. Nat Neurosci 12:535–540. doi:10.1038/nn.2303.Circular

Laureys S. 2005. The neural correlate of (un)awareness: Lessons from the vegetative state. Trends Cogn Sci 9:556–559. doi:10.1016/j.tics.2005.10.010

Mashour GA, Hudetz AG. 2017. Bottom-up and top-down mechanisms of general anesthetics modulate different dimensions of consciousness. Front Neural Circuits 11:1–6. doi:10.3389/fncir.2017.00044

Mumford JA, Davis T, Poldrack RA. 2014. The impact of study design on pattern estimation for single-trial multivariate pattern analysis. Neuroimage 103:130–138. doi:10.1016/j.neuroimage.2014.09.026

Mumford JA, Turner BO, Ashby FG, Poldrack RA. 2012. Deconvolving BOLD activation in event-related designs for multivoxel pattern classification analyses. Neuroimage 59:2636–2643. doi:10.1016/j.neuroimage.2011.08.076

Nagel T. 1974. What Is It Like to Be a Bat? Philos Rev 83:435–450. doi:10.4159/harvard.9780674594623.c15

Northoff G, Huang Z. 2017. How do the brain’s time and space mediate consciousness and its different dimensions? Temporo-spatial theory of consciousness (TTC). Neurosci Biobehav Rev 80:630–645. doi:10.1016/j.neubiorev.2017.07.013

O’Shea SM, Wong LC, Harrison NL. 2000. Propofol increases agonist efficacy at the GABA(A) receptor. Brain Res 852:344–348. doi:10.1016/S0006-8993(99)02151-4

Oosterhof NN, Connolly AC, Haxby J V. 2016. CoSMoMVPA: Multi-modal multivariate pattern analysis of neuroimaging data in matlab/GNU octave. Front Neuroinform 10:27. doi:10.3389/fninf.2016.00027

Overgaard M, Rote J, Mouridsen K, Ramsøy TZ. 2006. Is conscious perception gradual or dichotomous? A comparison of report methodologies during a visual task. Conscious Cogn 15:700–708. doi:10.1016/j.concog.2006.04.002

Pereira F, Mitchell T, Botvinick M. 2009. Machine learning classifiers and fMRI: a tutorial overview. Neuroimage 45:S199–S209. doi:10.1016/j.neuroimage.2008.11.007

Phillips WA, Bachmann T, Storm JF. 2018. Apical Function in Neocortical Pyramidal Cells: A Common Pathway by Which General Anesthetics Can Affect Mental State. Front Neural Circuits 12:1–15. doi:10.3389/fncir.2018.00050

Qiu M, Scheinost D, Ramani R, Constable RT. 2017. Multi-modal analysis of functional connectivity and cerebral blood flow reveals shared and unique effects of propofol in large-scale brain networks. Neuroimage 148:130–140. doi:10.1016/j.neuroimage.2016.12.080

Ratcliff R. 1993. Methods for dealing with response time outliers. Psychol Bull 114:510–532.

Sandberg K, Andersen LM, Overgaard M. 2014. Using multivariate decoding to go beyond contrastive analyses in consciousness research. Front Psychol 5:1–6. doi:10.3389/fpsyg.2014.01250

Sarasso S, Boly M, Napolitani M, Gosseries O, Charland-Verville V, Casarotto S, Rosanova M, Casali AG, Brichant JF, Boveroux P, Rex S, Tononi G, Laureys S, Massimini M. 2015. Consciousness and complexity during unresponsiveness induced by propofol, xenon, and ketamine. Curr Biol 25:3099–3105. doi:10.1016/j.cub.2015.10.014

Schrouff J, Perlbarg V, Boly M, Marrelec G, Boveroux P, Vanhaudenhuyse A, Bruno MA, Laureys S, Phillips C, Pélégrini-Issac M, Maquet P, Benali H. 2011. Brain functional integration decreases during propofol-induced loss of consciousness. Neuroimage 57:198–205. doi:10.1016/j.neuroimage.2011.04.020

Searle JR. 2000. Consciousness. Annu Rev Neurosci 23:557–578.

Sheikh UA, Carreiras M, Soto D. 2019. Decoding the meaning of unconsciously processed words using fMRI-based MVPA. Neuroimage 191:430–440. doi:10.1016/j.neuroimage.2019.02.010

Soto D, Sheikh UA, Rosenthal CR. 2019. A Novel Framework for Unconscious Processing. Trends Cogn Sci 23:372–376. doi:10.1016/j.tics.2019.03.002

Stelzer J, Chen Y, Turner R. 2013. Statistical inference and multiple testing correction in classification-based multi-voxel pattern analysis (MVPA): Random permutations and cluster size control. Neuroimage 65:69–82. doi:10.1016/j.neuroimage.2012.09.063

Suzuki M, Larkum ME. 2020. General Anesthesia Decouples Cortical Pyramidal Neurons. Cell 180:666–676.e13. doi:10.1016/j.cell.2020.01.024

Tononi G, Boly M, Massimini M, Koch C. 2016. Integrated information theory: From consciousness to its physical substrate. Nat Rev Neurosci 17:450–461. doi:10.1038/nrn.2016.44

Trapani G, Altomare C, Sanna E, Biggio G, Liso G. 2012. Propofol in Anesthesia. Mechanism of Action, Structure-Activity Relationships, and Drug Delivery. Curr Med Chem 7:249–271. doi:10.2174/0929867003375335

Tsuchiya N, Koch C. 2005. Continuous flash suppression reduces negative afterimages. Nat Neurosci 8:1096–1101. doi:10.1038/nn1500

Varoquaux G, Raamana PR, Engemann DA, Hoyos-Idrobo A, Schwartz Y, Thirion B. 2017. Assessing and tuning brain decoders: Cross-validation, caveats, and guidelines. Neuroimage 145:166–179. doi:10.1016/j.neuroimage.2016.10.038

Veselis RA, Feshchenko VA, Reinsel RA, Beattie B, Akhurst TJ. 2005. Propofol and thiopental do not interfere with regional cerebral blood flow response at sedative concentrations. Anesthesiology 102:26–34. doi:10.1097/00000542-200501000-00008

Wang L, Mruczek REB, Arcaro MJ, Kastner S. 2015. Probabilistic maps of visual topography in human cortex. Cereb Cortex 25:3911–3931. doi:10.1093/cercor/bhu277

Wislowska M, Del Giudice R, Lechinger J, Wielek T, Heib DPJ, Pitiot A, Pichler G, Michitsch G, Donis J, Schabus M. 2017. Night and day variations of sleep in patients with disorders of consciousness. Sci Rep 7:1–11. doi:10.1038/s41598-017-00323-4

