## Supplementary figures for "A reduced level of consciousness affects non-conscious processes"

**This PDF file includes:**

**Supplementary figures and captions 1 to 4**

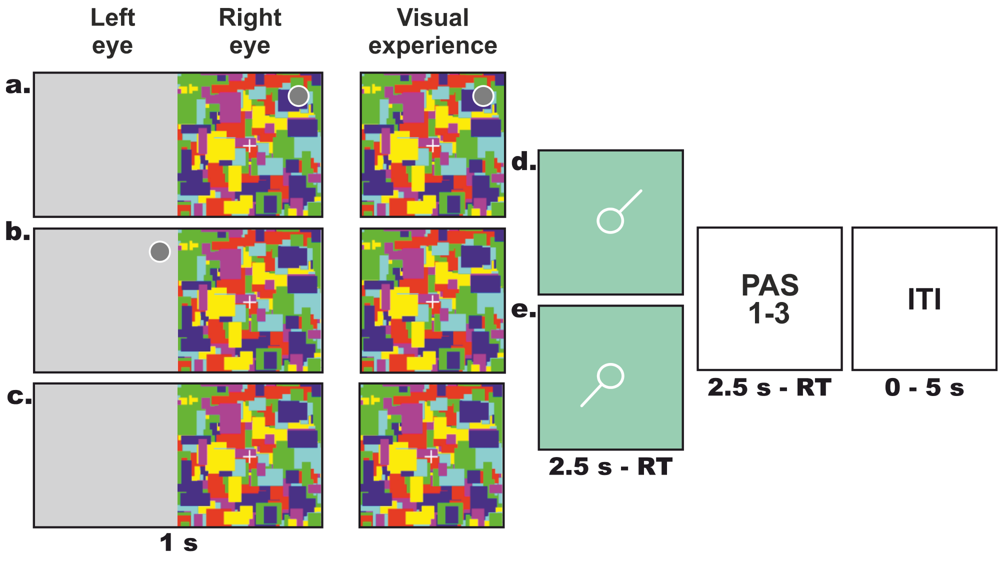


**Fig. S1: Visuospatial task using the CFS paradigm**. The task was composed of three presentation conditions. For each of the three conditions, Mondrians were displayed to the right eye. (**a.**) In the conscious condition, a full gray disc (sample presented at one of the 4 quadrants) was presented to the right eye (superimposed on the Mondrians). (**b.**) In the non-conscious condition, the sample was presented to the left eye, superimposed on a gray background and (**c.**) in the absent condition, only the gray background without the gray disc was presented to the left eye. The central column illustrates the visual experience of participants. After the 1 s sample presentation, a probe pointing to one of the 4 quadrants was presented for a maximum of 2.5 s or until participants decided whether it was pointing to the correct location of the disc (yes/no response). The probe could point (**d.**) to the correct location of the sample (match), or (**e.**) to an incorrect location (non-match). Participants then estimated, within 2.5 s, their conscious experience of the disc on a three-point perceptual awareness scale (PAS) from 1: “no visual experience” to 3: “clear visual experience”. Finally, trials were separated by an inter-trial interval (ITI). Note that for the purpose of illustration only, the gray dot is encircled in white. RT: response time.

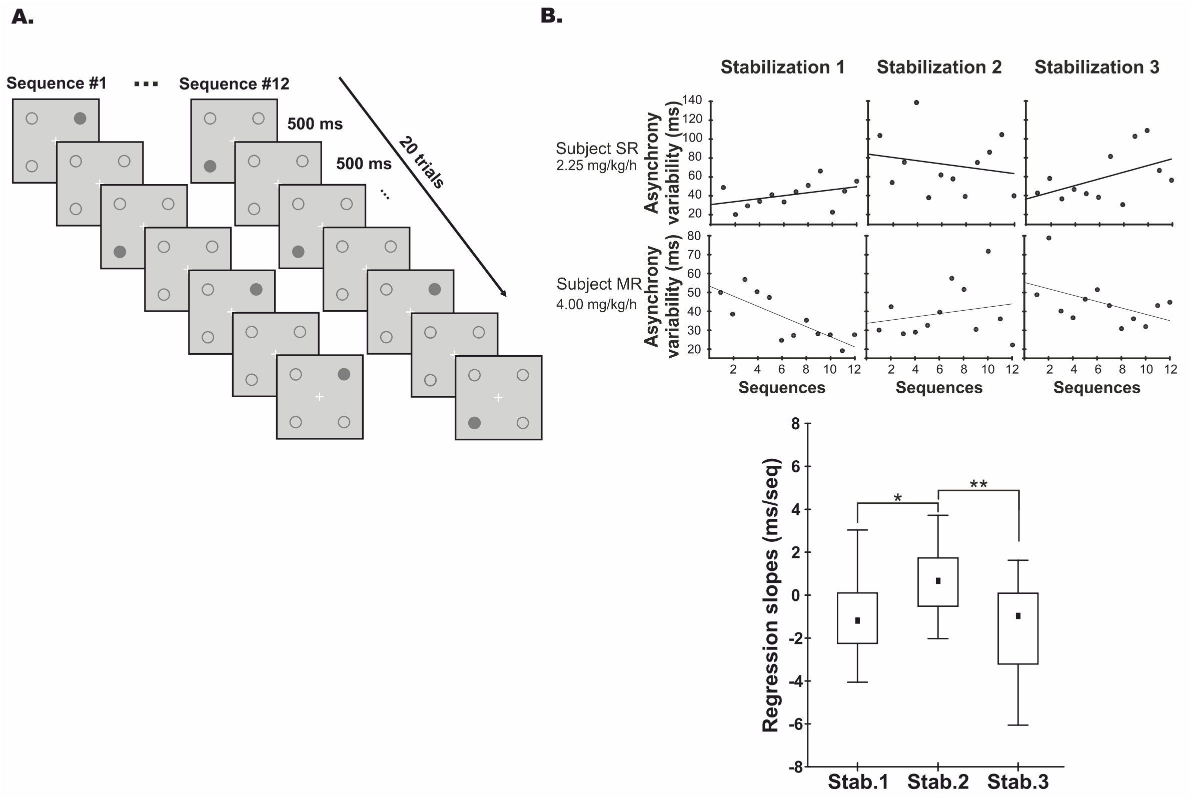


**Fig. S2: Illustration of procedure and performance for the visuo-motor "metronome task" performed during session 1 (pre-fMRI) and 3 (fMRI). A.** In the visual metronome task, the stimulus (full gray disc) displayed to the dominant-eye (right eye) was mainly presented (85% of the time) at one of the four quadrants followed by a 500 ms inter-trial interval, leading to a presentation of the stimulus with a tempo of 1 Hz. In total, 12 sequences composed of 3 sequences per location, each containing 20 trials, were presented in a pseudo-random order. **B. Estimation of the Propofol effect. (Top).** Illustration of performance for two representative participants, one starting with LS and the other starting with MS levels. The scatter plots represent the asynchrony variability (estimated with the standard deviation of the asynchrony) across the 12 sequences for the three stabilization periods where Propofol level was changed. The specific level of moderate sedation is indicated below the subject's initials. A positive slope indicating poor performance (increased variability) with the increase of Propofol reflects a decrease in arousal.

Correspondingly, a negative slope indicating better performance (reduced variability) with the decrease of Propofol indicates an increase in arousal. **(Bottom) Estimation of the Propofol effect at the group level.** The boxplots represent the average of the regression slopes for the three stabilization periods. Note that the first and third transitions are significantly different from the second one, confirming the correct manipulation of the sedation level before each block of the main visuospatial task. Stab. = Stabilization period; $*p < 0.05$; $**p < 0.01$.

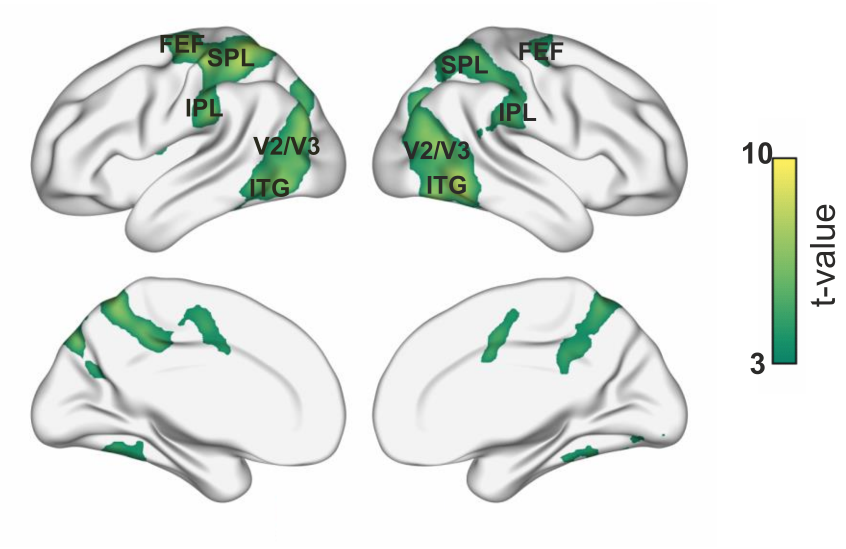


**Fig. S3: Conscious visuospatial network.** Activation map from the univariate analysis (Conscious > Absent), revealing a bilateral temporo-parieto-occipital network involved in visuospatial processing. The map is thresholded at $p < 0.05$ family-wise error rate corrected at the cluster level, and a cluster-defining threshold of $p < 0.001$. The color scale represents the t-value. FEF: frontal eye field, SPL: superior parietal lobule, IPL: inferior parietal lobule, V2/V3: secondary and tertiary visual areas, ITG: inferior temporal gyrus.

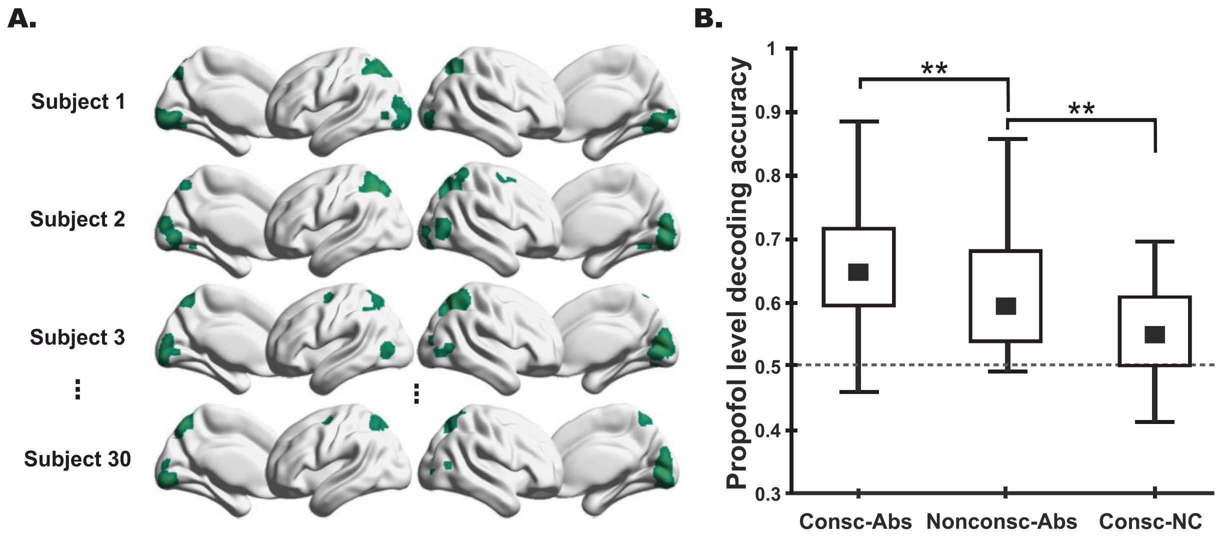


**Fig. S4: Effect of the sedation on conscious and non-conscious visuospatial processing. A.** Regions of interest (green) obtained with the maps of MVPA searchlight decoding the stimulus' side (right vs. left) in the metronome task. Note that to avoid any difference due to clusters size, we selected for each individual the same number of voxels as in the original ROI. **B.** Boxplots of decoding accuracy for the sedation level (low *vs.* moderate), for conscious and non-conscious conditions controlled for non-specific processing (-Abs), and for the conscious condition when controlling for non-conscious visuospatial processing (-NC). **$p < 0.01$.
